# Aging reveals domain-specific vulnerability to the chronic behavioral consequences of repetitive mild traumatic brain injury

**DOI:** 10.64898/2026.06.29.735325

**Authors:** Josh Karam, Julian Lopez, Leslie Ortiz, Aileen J. Anderson, Brian J. Cummings

**Affiliations:** Sue & Bill Gross Stem Cell Center, University of California-Irvine, Irvine, CA, 92697, United States of America; Institute for Memory Impairments & Neurological Disorders,University of California-Irvine, Irvine, CA, 92697, United States of America; Physical Medicine & Rehabilitation, University of California-Irvine, Irvine, CA, 92697, United States of America; Anatomy & Neurobiology, University of California-Irvine, Irvine, CA, 92697, United States of America

**Author notes:** Correspondence: Brian J. Cummings.

## Abstract

Older adults are among the fastest growing groups of traumatic brain injury (TBI) patients and sustain disproportionately poor chronic outcomes. Despite this, the preclinical aging-TBI literature is limited. Beyond the limited presence of aging TBI studies, most studies published in this domain use moderate-to-severe, open head models of TBI, rather than closed head models of mild TBI (mTBI) and repetitive mTBI (rmTBI), the most clinically prevalent presentation. Whether age modulates the chronic behavioral consequences of rmTBI is unknown. In the current study, young (3-4 months) and aged (18-19 months) male C57BL/6 mice received either five mTBIs on alternating days to model rmTBI or sham procedures and underwent behavioral testing in the chronic phase for spatial memory and anxiety-related behavior. Because cross-age behavioral comparisons are confounded by age-related declines in activity and by large sample sizes necessary to detection interaction effects, we applied a three-tier analytical framework combining within-age comparisons, sham-normalized inter-age comparisons, and factorial two-way ANOVA. Contrary to our hypothesis that aging would worsen rmTBI behavioral deficits, age produced domain-divergent effects. Spatial memory deficits were directionally consistent in both young and aged mice but was attenuated in the aged group. Conversely, anxiety-related behavior emerged selectively in the aged mice showing increased thigmotaxis. Locomotion was driven by age alone, with no injury effect, confirming that the aged anxiety signal was not a locomotor artifact. A post-hoc sensitivity analysis indicated that resolving the Age x Injury interaction effect would require at least 44 animals per group. These findings show that age shapes the affective, but not the cognitive, consequences of chronic rmTBI, and underscoring that statistical strategy is inseparable from design in factorial injury studies.

## Introduction

Traumatic brain injuries (TBI) are the <u>seventh</u> leading cause of disability globally, with ∼49 million prevalent cases and a collective ∼7 million years lived with disability1. Approximately 80% of TBI patients are diagnosed with mild TBI (mTBI), which can be concussive or sub-concussive2. A single mTBI typically has a favorable prognosis with full recovery within 3 months2. Approximately 15 percent of patients experience post-concussion symptoms which are exacerbated by repetitive mTBI (rmTBI)2. Further, adults aged 65 and older are among the fastest growing groups of TBI patients, sustaining disproportionately poor functional outcomes that persist months after injury, largely independent of initial injury severity3–6. TBI sustained in older age carries elevated long-term risk of dementia and progressive cognitive decline4,5, thus making the aged brain a central, and understudied, problem in chronic management of TBI.

Despite the presence of a clinical need to study the age-specific sequelae of TBI, the literature remains limited and conflicting. For example, Ritzel *et al.,* found that aged mice (18 months old) exhibited worse anxiety-related, motor, and locomotor outcomes compared to young mice (3 months old) acutely after injury (<7 days post-injury) using a controlled cortical impact (CCI) model of TBI7. However, Islam *et al.,* in the same mouse strain, same age ranges, and similar CCI model reported diferential behavioral outcomes where young mice exhibited more disinhibition and aged mice exhibited more anxiety-related behaviors at 30 days post-injury. Additionally, more severe neuropathology was reported in young mice compared to aged mice8. The gap in these findings exemplify that the need to study age-specific efects of TBI, as age does not simple worsen TBI, but perhaps drives domain-specific and region-specific injury responses. Even though aging studies in the TBI field are increasing, the majority of these studies focus on moderate-to-severe, open head models of TBI such as CCI or fluid percussion injuries7–12, presenting a clinically consequential gap: age-specific efects after rmTBI12. Studies involving rmTBI are imperative as rmTBI is the dominant clinical presentation, especially in aged populations, aging veterans, and former contact-sports participants. Whether age modulates the chronic behavioral consequences of repetitive mild traumatic brain injuries is unknown.

Here, we specifically test whether age alters chronic behavioral outcomes after rmTBI. Young (3-4 months) and aged (18-19 months) male C57BL/6 mice received five mTBIs on alternating days or sham procedures and underwent testing for spatial memory (Barnes Maze) and anxiety-related behavior (Elevated Plus Maze and Open Field Test) in the chronic phase of injury (**Fig 1**). There are two analytical confounds present in any aging TBI study with behavioral outcomes. First, mice become less active and exploratory as they age which confounds their performance in many behavioral tasks and any cross-age comparison based those performances13. Second, detecting an Age x Injury interaction afect requires substantially larger group sizes than detecting a comparable main efect (either age or injury alone) which is often infeasible for a single research lab14,15. On the other hand, inferring age-dependence from contrasting significance in independent groups is also a statistical error16. To address both of these constraints, we implemented a three-tier framework combining within-age comparisons, normalized inter-age comparisons, and factorial two-way ANOVA. We hypothesized that aging would cause worse spatial memory deficits and anxiety-related behavior. Interestingly, across all three tiers of our analysis, age produced domain-divergent chronic outcomes after rmTBI: spatial memory deficits in both young and aged mice, anxiety-like behavior only in aged mice, and locomotion preserved in both young and aged animals.

**Figure 1:**
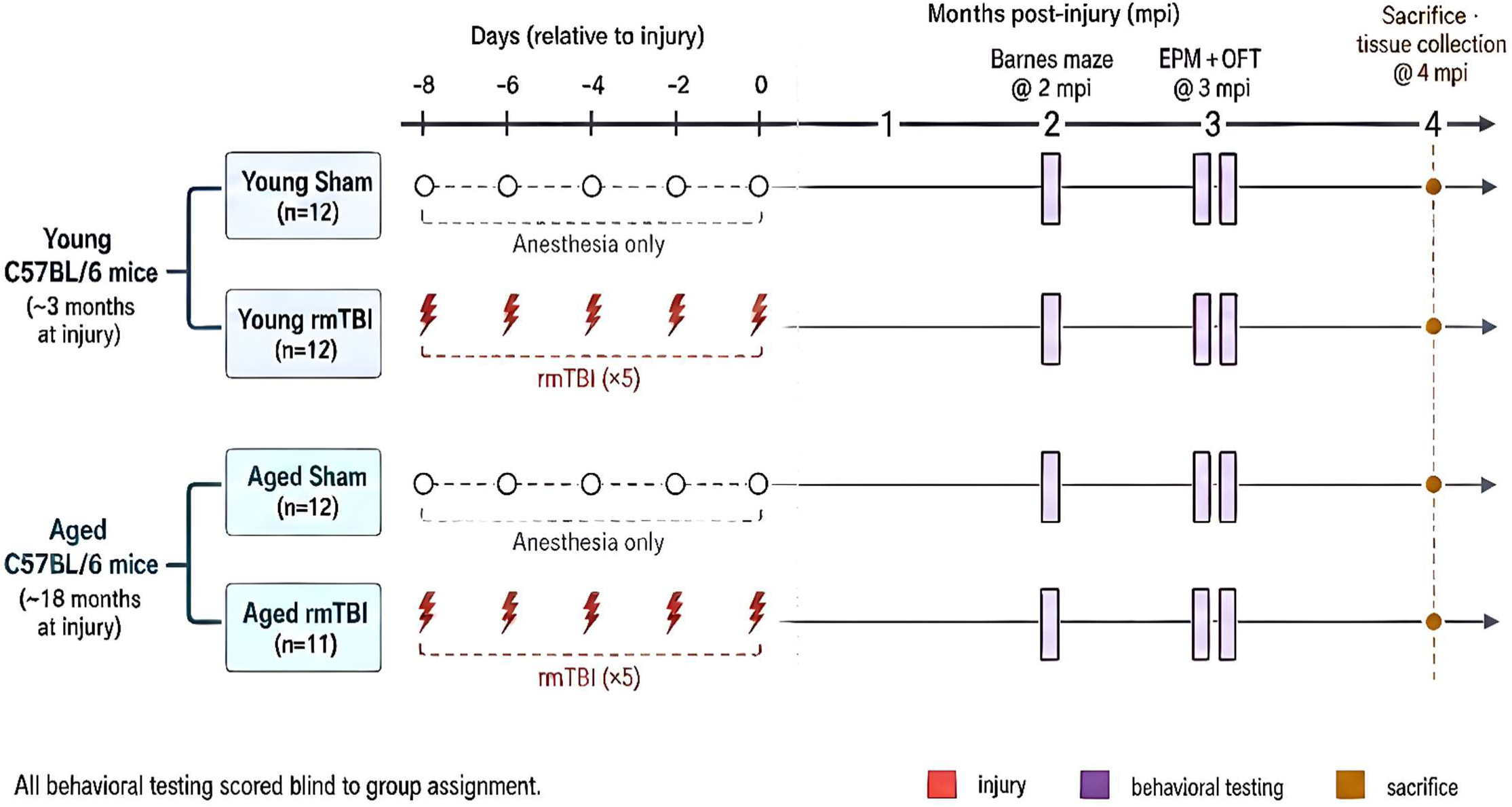
Experimental Timeline. Young (3-4 months) and aged (18-19 months) male C57BL/6 mice received five mTBIs on alternating days or sham procedures and underwent testing for spatial memory (Barnes Maze) at 2 mpi and anxiety-related behavior (Elevated Plus Maze and Open Field Test) at 3 mpi. Experimental timeline schematic was AI-generated using FigureLabs.

## Methods

### Animals

All experiments in the following study were approved by the Institute for Animal Care and Use Committee at the University of California, Irvine, and were carried out per the Guide for the Care and Use of Laboratory Animals. Male C57BL/6 mice were used in this study. Young (2-3 months old, n=12/group) and aged (18-19 months old, n=11-12/group) mice were acclimated for at least one week prior to being injured. All animals were housed in a controlled humidity and temperature environment with free access to food and water, as well as access to houses and nestlets for enrichment. Mice were kept on a 12 hour (6:30 am-6:30 pm) light/dark cycle.

### Repetitive mild traumatic brain injury model (rmTBI)

Prior to injury, mice were randomly assigned to experimental groups. Experimenters remained blinded through the entirety of the study. Animals were given rmTBI as previously described^17–19^. Briefly, mice were anesthetized using 2.5% isoflurane. A pneumatic controlled cortical impact device (TBI-0310 Head Impactor, Precision Systems and Instrumentation, LLC, Fairfax Station, VA) was used to deliver mTBI. All mice spent 7 min under anesthesia, including shams. The mouse was placed onto modified Marmarou foam and positioned to breathe 2.5% isoflurane from a nose cone. A 5-mm diameter probe tip was used to deliver an impact with speed 5.0 m/s, 1.0 mm depth, and 50 ms dwell time. In total, mice were under anesthesia for 7 min from knockdown to mTBI impact. The isoflurane percentages were designed to minimize head movement from respiration while zeroing the piston and during impact while keeping the animal unresponsive to a toe pinch reflex. Following impact, animals were moved to a recovery area where righting time was recorded. Sham animals underwent the same 7-min procedure with anesthesia but were not impacted. This procedure was repeated every other day for 9 days to accumulate 5 mTBIs to model rmTBI (**Fig 1**).

### Barnes Maze

Barnes Maze was used to test spatial learning and memory in mice at 2 months post-injury (mpi). A 20-hole Barnes Maze apparatus was placed in a room with Noldus Behavioral Testing video acquisition and analysis software. Barnes Maze testing was performed in ambient lighting with no white noise or other distressing factors. Mice were transferred to the behavioral testing room at least one hour before testing began. On Day 1, all mice underwent a habituation trial for 5 minutes where they were placed in the center of the maze and allowed to explore for 5 minutes or until they found the target escape box. At the end of the habituation trial, all mice were placed in the target escape box for 90 seconds. On Days 2-5, mice underwent three acquisition trials each day (9 am–11 am, 12 pm-2 pm, and 3 pm-5 pm) where they were placed on the maze for 3 minutes or until they found the target escape box. If in the trial, they did not find the target escape box, they were placed inside it for 30 seconds before being returned to their home cage. The target escape box was moved after habituation to a different location for the acquisition where it remained for all of acquisition. On Day 6, the mice underwent the short-term memory probe trial, 24 hours after the last acquisition trial. The target escape box was removed and mice were placed in the center of the maze and allowed to explore for 90 seconds. Video recordings were tracked and analyzed using Noldus Ethovision XT 14 and exported for statistical analysis in R. During the probe trial, Animal 13 (Young Sham group) did not move at all, and therefore was excluded from statistical testing due to not performing the test. Spatial memory domain scoring was computed by z-scoring all primary and secondary metrics within age groups from the Barnes Maze probe trial with directionality chosen so that a higher z-score correlates with worse spatial memory. The z-scores for each metric were then averaged to assign a spatial memory domain score for each animal^20–22^.

### Elevated Plus Maze (EPM)

One method of assessing anxiety was by comparing time spent in the open arms and closed arms, as well as the number of entries into either of an elevated plus maze apparatus (Noldus Information Technology, Leesburg, VA) as previously described^17,23,24^. Animals are placed at the center point of a maze covered by a cup for 30 seconds. After 30 seconds, mice are released to explore the EPM for 5 minutes. EPM was performed in a dark room with red overhead lighting, near-infrared illumination under the maze arms for contrast and camera detection, and 57 dB white noise. For anxiety domain scoring, all four metrics were z-scored with directionality chosen so that a higher z-score correlates with greater anxiety-related behavior then averaged. The EPM average z-score was averaged with the z-score for Open Field Test to compute the anxiety domain score.

### Open Field Test (OFT)

The other anxiety-related behavioral task used in the is study was the Open Field Test (OFT). OFT was performed in the UCI Behavioral Testing Core in a dimly lit room with no white noise. Mice were placed in the OFT apparatus and allowed to roam freely for 10 minutes. Mice were recorded and tracked using Noldus Ethovision XT 18 to calculate the percentage of time spent in the outer zone (thigmotaxis) and inner zone, as well as total distance traveled. Time spent in each zone was z-scored as for EPM and averaged for the OFT test. The averaged OFT z-score was averaged with the EPM z-score to compute the anxiety domain score.

### Statistical Analysis

Comparisons between young and aged animals’ behavioral outcomes are confounded by two pre-existing analytical constraints. First, it is well-established that as animals age, they become less active and exploratory^7,13^ which can confound performances in the behavioral tests in this study complicating direct cross-age comparison of injury effects. Additionally, it is established in the field of behavioral statistics that detecting an interaction effect (age x injury) requires a significantly larger sample size than that needed for an equivalent main effect^14–16^. This necessary increase in sample size presents serious logistical challenges for single-laboratory rodent studies that are infeasible. To account for these two main confounds in this study, we used a pre-specified three-tiered statistical analysis approach.

#### Tier 1: Within-age comparisons

Tier 1 analysis is designed to preserve sensitivity of injury effects within age groups independent of age-related activity differences. First, we tested normality of the data in each group via Shapiro-Wilk normality test. If both groups passed, then Welch’s two-sample t-test was used for hypothesis testing. If at least one group failed, then Mann-Whitney U test was used for hypothesis testing. It has been shown consistently in the literature that TBI impairs spatial memory which our lab has confirmed in multiple studies^12,17,23–27^. Due to this, we had an *a priori* hypothesis that mice with rmTBI would have worse performance on the Barnes Maze task and therefore employed a one-tailed test for hypothesis testing. For EPM and OFT, there has been inconsistency in the literature regarding the effect of rmTBI on EPM performance in rodents^17,19,28^, so we utilized two-tailed hypothesis testing. Beyond hypothesis testing, we also calculated effect sizes using either Cohen’s d (parametric) or rank-biserial r (non-parametric.

#### Tier 2: Normalized inter-age comparison

Tier 2 analysis is designed to test whether the magnitude of the injury response, relative to each cohort’s own baseline, differs between ages. Each rmTBI animal’s outcomes are expressed as a percentage of the within-age Sham mean. We performed either one-sample t tests (parametric) or Wilcoxon signed-ranks (non-parametric) against 100 for each cohort. After normalization, either a Welch’s t test or a Mann-Whitney U test was used for comparison of Young normalized to Aged normalized distributions. Spatial Memory Domain and Anxiety Domain scores were excluded from this analysis since they are z-scores and normalizing a standardized z-score to a near-zero reference mean is not interpretable.

#### Tier 3: Two-way ANOVA

Tier 3 analysis formally deconvolves age and injury main effects within the correct factorial framework. Interaction testing here is explicitly exploratory due to the underpowered sample sizes in this study. Effect size is calculated using partial ω²p (partial omega-squared). Post-hoc multiple comparisons was performed to test within age and within injury contrasts using Holm-Bonferroni correction. A post-hoc sensitivity analysis was performed using the data in this study to determine the statistical power of this study and the appropriate sample size to detect an interaction effect.

## Results

### RmTBI impairs spatial memory in both young and aged mice

Both the young and aged cohorts exhibited spatial memory deficits, though to different extents, across multiple Barnes Maze probe metrics. Young mice with rmTBI showed significant impairment of spatial memory across multiple Barnes Maze metrics, while aged mice with rmTBI only showed mild impairment with multiple metrics trending towards significance with modest effect sizes.

In our Tier 1 analysis of the young cohort of mice, rmTBI induced significant increases in latency to target (Mann-Whitney right-tail, *p*=0.0363, *r*=0.44, medium, **Fig 2C**) and percentage of time spent in the opposite quadrant (Mann-Whitney right-tail, *p*=0.0105, *r*=0.58, large, **Fig 2G**). Conversely, rmTBI also induced a significant decrease in percentage of time spent in the target quadrant (Welch t left-tail, *p*=0.0363, *d*= −0.80, medium, **Fig 2E**). Furthermore, rmTBI mice showed a trending increase in primary path length (Mann-Whitney right-tail, *p*=0.0658, *r*=0.38, medium, **Fig 2A**). Spatial memory domain composite scoring was used to assign an overall score that accounts for all metrics from the Barnes Maze where a higher score is correlated with worse spatial memory. Mice with rmTBI had significantly higher spatial memory domain scores (Welch t right-tail, *p*=0.0174, *d=*0.94, large, **Fig 2I**).

**Figure 2:**
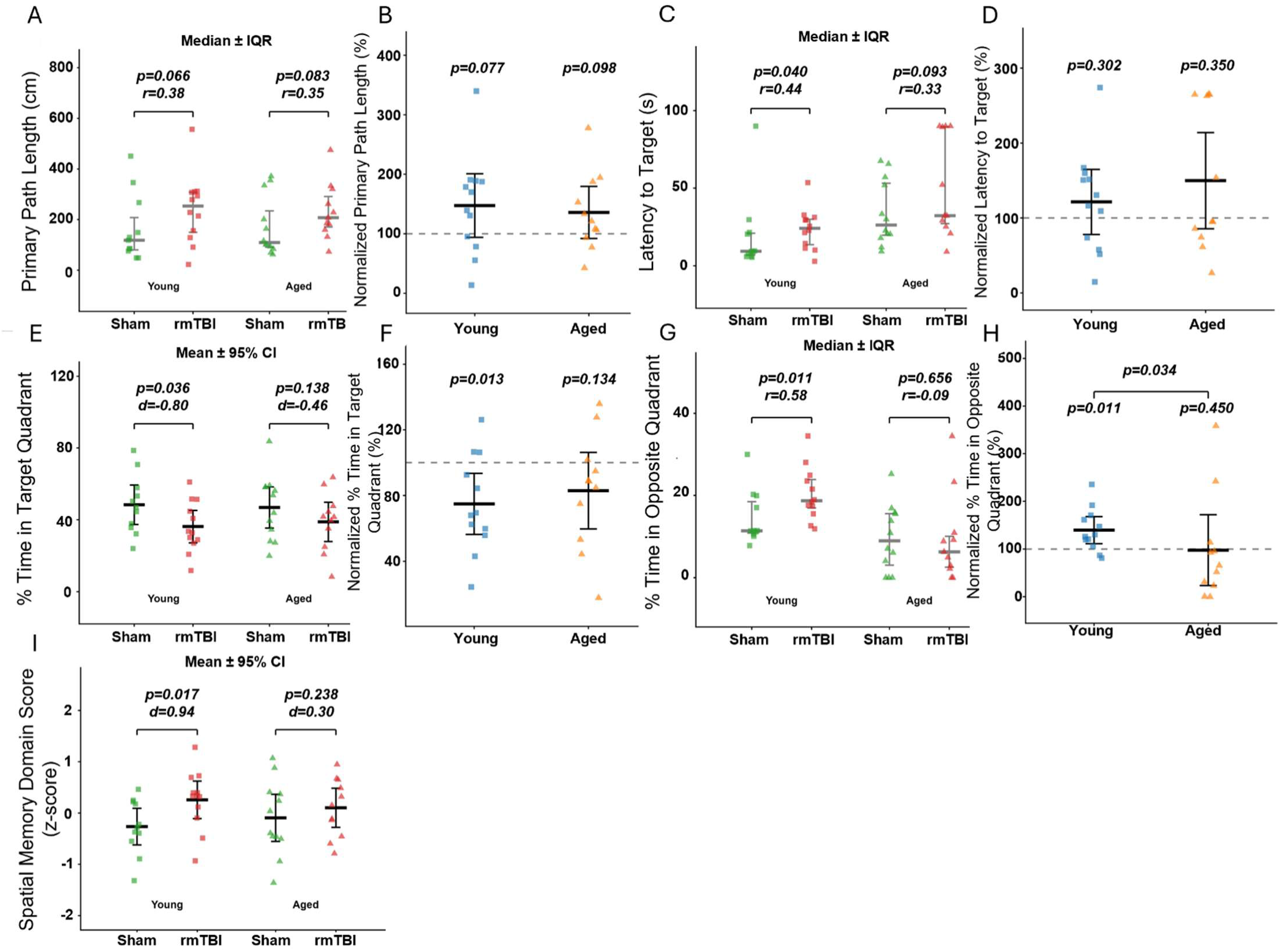
RmTBI induces spatial memory deficits in young and aged mice at 2 mpi. Both the young and aged cohorts were tested for spatial memory retention at 2 mpi. 24 hour probe trial outcomes are reported for both cohorts using the following primary metrics: Primary Path Length (A-B), Latency to Target (C-D), % Time in Target Quadrant (E-F), and % Time in Opposite Quadrant (G-H). Tier 1 within-age comparisons (A, C, E, G) and Tier 2 normalized inter-age comparisons (B, D, F, H) are shown. Barnes Maze primary metrics and secondary metrics were z-scored to give an overall spatial memory domain score (I). Dependent on normality, either the mean ± 95% confidence interval or the median ± interquartile range is shown. p-values and effect sizes are shown.

Our Tier 1 analysis of the aged cohort of mice showed rmTBI did not significantly alter any of the Barnes Maze metrics. However, there were trending increases in primary path length (Mann-Whitney right-tail, *p*=0.0831, *r*=0.35, medium, **Fig 2A**) and latency to target (Mann-Whitney right-tail, *p*=0.0927, *r*=0.33, medium, **Fig 2C**). There were no significant effects on percentage of time spent in the target quadrant (Welch t left-tail, *p*=0.1380, *d=* −0.46, small, **Fig 2E**) or opposite quadrant ((Mann-Whitney right-tail, *p*=0.6562, *r*= −0.09, small, **Fig 2G**), as well as spatial memory domain score (Welch t right-tail, *p*=0.2379, **Fig 2I**).

Comparing the within-age sham normalized rmTBI groups between ages in our Tier 2 analysis shows overall convergence between ages across the primary Barnes Maze metrics of primary path length (Welch t, *p*=0.7151, *d*=0.15, small, **Fig 2B**), latency to target (Mann-Whitney, *p*=0.6888, *r*= −0.11, small, **Fig 2D**), and percentage time spent in target quadrant (Welch, *p*=0.5575, *d*= −0.25, small, **Fig 2F**). Interestingly, our inter-age analysis showed that young rmTBI mice spent significantly less time in the opposite quadrant (Mann-Whitney, *p*=0.0337, *r*=0.53, large, **Fig 2H**) compared to aged rmTBI mice.

While the within-age analysis revealed Barnes Maze metrics showing impaired spatial memory in young mice and to a lesser extent aged mice, our Tier 3 analysis (**Table 1**) confirms our initial expectation that there would not be an observed age x injury interaction effect, primarily due to the dominance of age as a significant main effect in the primary metrics for latency to target (*p=*0.0056, ω²p=0.138, medium) and percentage of time spent in the opposite quadrant (*p*=0.0014, ω²p=0.187, large). Interestingly, injury is a significant main effect in percentage of time spent in target quadrant (*p*=0.0419, ω²p=0.071, medium), and a trending main effect in primary path length (*p*=0.0675, ω²p=0.054, small) and spatial memory domain score (*p*=0.0515, ω²p=0.063, medium). Post-hoc multiple comparisons show the young rmTBI have a significantly longer latency to target (*p*=0.0343, *d*=1.15, large) and spend significantly more time in the opposite quadrant (*p*=0.0083, *d*= −1.37, large).

**Table 1:** Tier 3 Factorial Analysis Results. Two-way ANOVA with a post-hoc Holm corrected multiple comparisons test was performed for all three behavioral tasks. Here, we report *p*-values and effect sizes for the main effects, interaction, and post-hoc multiple comparisons.

| Metric | Age main effect | Injury main effect | Age × Injury interaction | Young: Sham vs rmTBI | Aged: Sham vs rmTBI | Shams: Young vs Aged | rmTBI: Young vs Aged |
| --- | --- | --- | --- | --- | --- | --- | --- |
| Primary path length (cm) | 0.9129<br>$\omega^2_p = 0.000$ (neg.) | 0.0675<br>$\omega^2_p = 0.054$ (small) | 0.8259<br>$\omega^2_p = 0.000$ (neg.) | 0.5813<br>$d = 0.62$ | 0.7449<br>$d = 0.49$ | 1.0000<br>$d = 0.06$ | 1.0000<br>$d = -0.07$ |
| Latency to target (s) | 0.0056<br>$\omega^2_p = 0.138$ (medium) | 0.1375<br>$\omega^2_p = 0.024$ (small) | 0.3656<br>$\omega^2_p = 0.000$ (neg.) | 0.6739<br>$d = 0.18$ | 0.2799<br>$d = 0.72$ | 0.3013<br>$d = 0.61$ | 0.0343<br>$d = 1.15$ |
| % time in target quadrant | 0.8383<br>$\omega^2_p = 0.000$ (neg.) | 0.0419<br>$\omega^2_p = 0.071$ (medium) | 0.6683<br>$\omega^2_p = 0.000$ (neg.) | 0.3233<br>$d = -0.75$ | 0.7355<br>$d = -0.49$ | 1.0000<br>$d = -0.09$ | 1.0000<br>$d = 0.16$ |
| % time in opposite quadrant | 0.0014<br>$\omega^2_p = 0.187$ (large) | 0.2511<br>$\omega^2_p = 0.006$ (neg.) | 0.2131<br>$\omega^2_p = 0.010$ (small) | 0.2801<br>$d = 0.72$ | 0.9437<br>$d = -0.03$ | 0.2845<br>$d = -0.62$ | 0.0083<br>$d = -1.37$ |
| Spatial Memory Domain (z-score) | 0.9676<br>$\omega^2_p = 0.000$ (neg.) | 0.0515<br>$\omega^2_p = 0.063$ (medium) | 0.3687<br>$\omega^2_p = 0.000$ (neg.) | 0.1826<br>$d = 0.86$ | 1.0000<br>$d = 0.32$ | 1.0000<br>$d = 0.28$ | 1.0000<br>$d = -0.25$ |
| % time in open arms | 0.0050<br>$\omega^2_p = 0.136$ (medium) | 0.3127<br>$\omega^2_p = 0.001$ (neg.) | 0.0824<br>$\omega^2_p = 0.038$ (small) | 0.7580<br>$d = 0.21$ | 0.1604<br>$d = -0.83$ | 0.7580<br>$d = -0.36$ | 0.0066<br>$d = -1.40$ |
| % time in closed arms | 0.0497<br>$\omega^2_p = 0.060$ (small) | 0.8989<br>$\omega^2_p = 0.000$ (neg.) | 0.0779<br>$\omega^2_p = 0.044$ (small) | 0.5238<br>$d = -0.48$ | 0.5238<br>$d = 0.58$ | 0.8557<br>$d = 0.07$ | 0.0390<br>$d = 1.13$ |
| % time in outer zone | 0.0000<br>$\omega^2_p = 0.337$ (large) | 0.0207<br>$\omega^2_p = 0.062$ (medium) | 0.4013<br>$\omega^2_p = 0.000$ (neg.) | 0.2667<br>$d = 0.46$ | 0.0545<br>$d = 0.95$ | 0.0085<br>$d = 1.29$ | 0.0004<br>$d = 1.79$ |
| % time in inner zone | 0.0000<br>$\omega^2_p = 0.306$ (large) | 0.0367<br>$\omega^2_p = 0.051$ (small) | 0.6630<br>$\omega^2_p = 0.000$ (neg.) | 0.2235<br>$d = -0.50$ | 0.1509<br>$d = -0.76$ | 0.0088<br>$d = -1.29$ | 0.0024<br>$d = -1.54$ |
| Anxiety Domain (z-score) [6 metrics] | 1.0000<br>$\omega^2_p = 0.000$ (neg.) | 0.1171<br>$\omega^2_p = 0.033$ (small) | 0.5650<br>$\omega^2_p = 0.000$ (neg.) | 1.0000<br>$d = 0.30$ | 0.5306<br>$d = 0.64$ | 1.0000<br>$d = -0.16$ | 1.0000<br>$d = 0.18$ |

### RmTBI is anxiogenic in aged mice, but not young mice

Unlike in the Barnes Maze where both young and aged mice had similar spatial memory phenotypes, anxiety-related behaviors is exclusively altered in Aged rmTBI mice in both Elevated Plus Maze and Open Field Test. Young rmTBI mice showed no anxiety signal in either task for any metric.

Our Tier 1 analysis testing the effect of rmTBI within each age group showed that aged rmTBI mice exhibited much more anxiety-like behavior compared to aged sham. Both the OFT and EPM produced significant effect sizes. In EPM, aged rmTBI mice spend significantly less time in the open arms (Welch t, *p*=0.0265, *d*= − 1.02, large, **Fig 3C**) compared to aged sham mice, and there was no significant difference in time spent in the closed arms even though there is a medium effect (Welch t, *p*=0.1224, *d*=0.68, medium, **Fig 3A**). A similar phenomenon happens in the OFT, aged rmTBI mice spent significantly more time in the outer zone (Welch t, *p*=0.0468, *d*=0.89, **Fig 3E**) compared to aged sham mice. Interestingly, time spent in the inner zone was not significantly different, although there was a medium effect size (Welch t, *p*=0.1064, *d*= −0.72, medium, **Fig 3G**). An anxiety domain score was computed to assign an overall score for anxiety-related behavior across both tests where a higher anxiety domain score means higher anxiety-related behavior. Aged rmTBI mice had a significantly higher anxiety domain score compared to aged sham mice (Welch t, *p*=0.0312, *d*=1.01, large, **Fig 3I**). Young rmTBI mice and young sham mice had no detectable differences.

**Figure 3:**
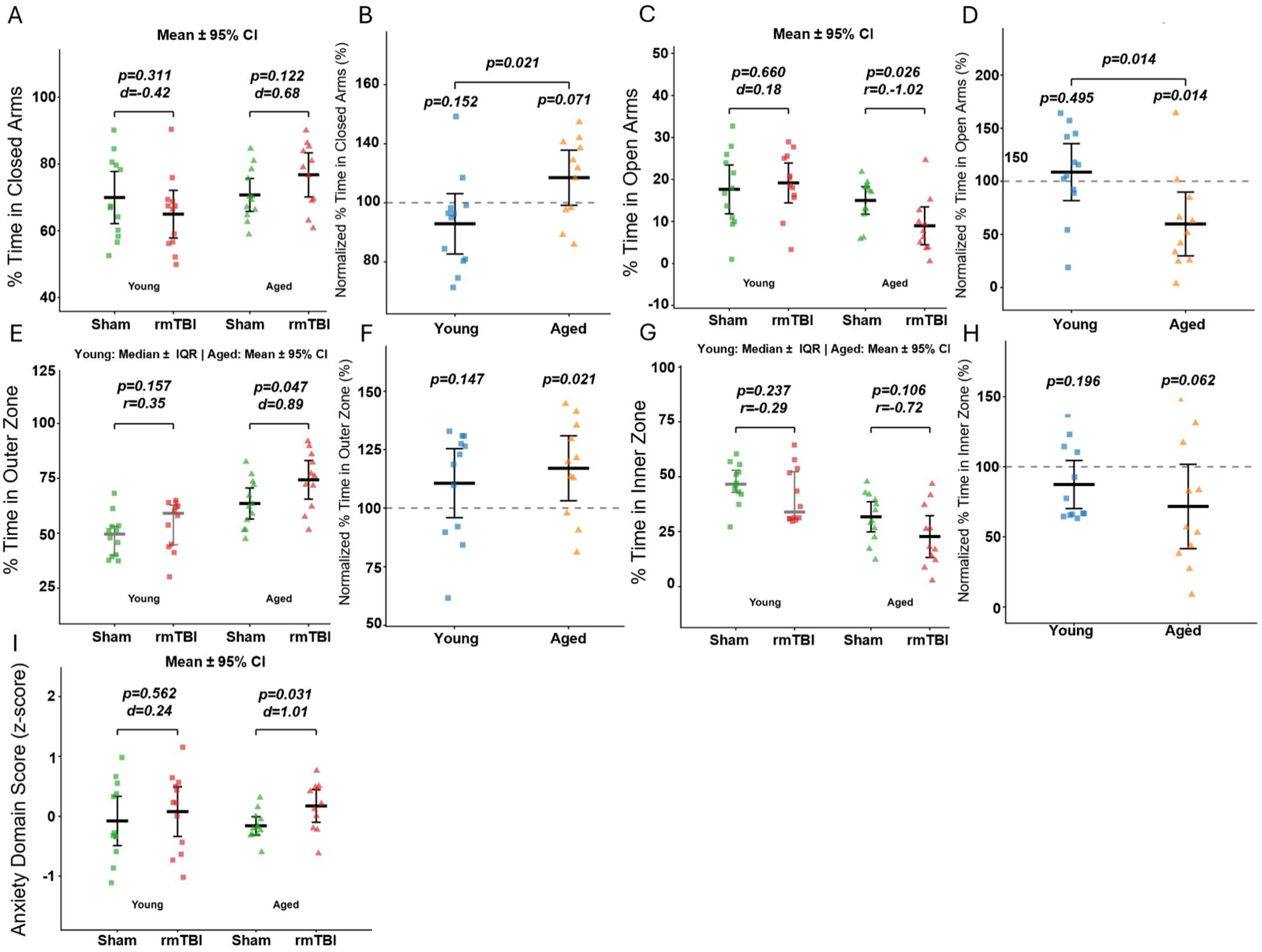
Anxiety is elevated in an age-dependent manner after rmTBI at 3 mpi. Both the young and aged cohorts were tested for anxiety-related behavior at 3 mpi. EPM metrics of % time in closed arms (A-B) and in open arms (C-D) are reported. OFT metrics of % time in outer zone (E-F) and in inner zone (G-H) are also reported. Tier 1 within-age comparisons (A, C, E, G) and Tier 2 normalized inter-age comparisons (B, D, F, H) are shown. EPM and OFT metrics were z-scored to give an overall anxiety domain score (I). Dependent on normality, either the mean ± 95% confidence interval or the median ± interquartile range is shown. p-values and effect sizes are shown.

Interestingly, inter-age comparisons of sham-normalized rmTBI groups in our Tier 2 analysis revealed that the aged mice exhibit more anxiety-related behavior compared to young mice in EPM, but not OFT. The sham-normalized aged rmTBI group spent significantly more time in the closed arms (Welch t, *p*=0.0208, *d*= − 1.04, large, **Fig 3B**) and significantly less time in the open arms (Welch t, *p*=0.0140, *d*=1.12, large, **Fig 3D**) compared to the sham-normalized young rmTBI group.

In our factorial two-way ANOVA analysis in Tier 3 (**Table 1**), there is still no significant interaction effect. However, the interaction effect is trending in the EPM metrics of percentage of time spent in closed arms (*p*=0.0779, ω²p=0.044, small) and open arms (*p*=0.0824, ω²p=0.038, small). Further, there is a significant injury effect in the OFT metrics of percentage of time spent in the outer zone (*p*=0.0207, ω²p=0.062, medium) and inner zone (*p*=0.0367, ω²p=0.051, small). Age is a significant main effect in percentage of time spent in closed arms (*p*=0.0497, ω²p=0.060, small), open arms (*p*=0.0050, ω²p=0.136, medium), outer zone (*p*<0.0001, ω²p=0.337, large), and inner zone (*p*<0.0001, ω²p=0.306, large). Age dominates post-hoc multiple comparisons with significant differences between young and aged mice in both sham groups and rmTBI groups. However, our post-hoc multiple comparisons reveal that aged rmTBI mice are trending towards spending more time in the outer zone of the OFT (*p*=0.0545, *d*=0.95, large) compared to aged sham mice. Interestingly, anxiety domain scores revealed no significant main effect, interaction effect, or post-hoc multiple comparisons.

### Locomotor activity in behavioral tasks is driven by age validating three-tier statistical rationale

Based on previous literature showing significant changes in baseline locomotor and exploratory activity as mice age^7,13^, we hypothesized that baseline activity differences would confound our age x injury analysis in these behavioral tasks where mice need to move and explore. In our Tier 1 within-age analysis, rmTBI had no significant effect on total distance traveled in Barnes Maze, EPM, or OFT. The lack of an injury effect in total distance for EPM and OFT supports that reduced open arm occupancy and increased outer zone time are specific anxiogenic signals and not artifacts of reduced locomotor capacity. Interestingly, while rmTBI did not significantly increase total distance traveled in young mice, it did produce a medium effect (Welch t, *p*=0.2486, *d*=0.52, medium). In our Tier 2 analysis, sham-normalized young rmTBI mice showed a significant increase in distance traveled against 100% (one-sample t, *p*=0.0166, *dz*=0.81, large). Not surprisingly, our Tier 3 analysis revealed age as a significant main effect for total distance traveled in Barnes Maze (*p*<0.0001, ω²p=0.431, large), OFT (*p*=0.0227, ω²p=0.090, medium), and EPM (*p*=0.0019, ω²p=0.177, large). Further, age is also a significant main effect for Barnes Maze and EPM metrics that are not directly related to distance traveled but can be confounded by it. For Barnes Maze, age was a significant main effect for errors made before target ((*p*=0.0007, ω²p=0.211, large) and total errors (*p*<0.0001, ω²p=0.391, large) with significant post-hoc multiple comparisons across age groups in both shams and rmTBI. For EPM, age was a significant main effect for the number of closed arm entries (*p*=0.0052, ω²p=0.144, large) and open arm entries (*p*=0.0025, ω²p=0.168, large) with significant post-hoc multiple comparisons across age groups in both shams and rmTBI. Interestingly, these metrics showed no significant injury effect across in our Tier 1 analysis.

## Discussion

This study shows that aging drives domain divergence in chronic behavioral outcomes after rmTBI. Spatial memory is affected to varying extents in both young and aged mice after rmTBI. However, anxiety-related behavior is selectively elevated in aged mice, which is confirmed by a lack of injury effect on the distance traveled in anxiety-related behavioral tests. All three tiers of analysis converge on this finding.

We have previously shown that our model of rmTBI induces chronic cognitive deficits in spatial memory in the Morris Water Maze task in young mice at 2- and 6- mpi^17^. While Morris Water Maze and Barnes Maze both test spatial memory, swimming in Morris Water Maze is known to induce neurochemical changes affecting spatial learning and memory^29^. Further, it has been shown that C57BL/6 mice perform better in Barnes Maze than Morris Water Maze^30^. For this study, we transitioned to Barnes Maze to test spatial memory deficits in our rmTBI model and expanded our study to include aged animals. Our findings reproduce our past results showing spatial memory impairment in young mice after rmTBI, using a distinct hippocampal-dependent task. Beyond replication, we extend our findings to aged animals showing a directionally consistent, but attenuated effect size in the aged cohort (spatial memory domain *d=*0.30 in aged vs *d*=0.94 in young, **Fig 2I**). We interpret this reduced effect size as the baseline age-related spatial impairment compressing dynamic range for a detectable effect. Our Tier 2 normalization analysis to compare the rmTBI response between ages supports this interpretation as the sham-normalized rmTBI groups in each age are statistically indistinguishable (**Fig 2B-H**). Beyond that, our Tier 3 two-way ANOVA (**Table 1**) shows the injury main effect to be near-significant with a medium effect size (*p*=0.0515, ω²p=0.063) precisely because spatial memory is impaired in both age cohorts. Placed in the context of the literature, our findings address a critical gap. Many studies have reported acute to chronic memory deficits after rmTBI^17,31–34^, however, there is a dearth of literature examining how age factors into outcomes after rmTBI^12,35^. In more severe models of TBI, such as controlled cortical impact (CCI) and fluidic percussion injury (FPI), there has been an slight increase in recent literature showing that aged rodents exhibit similar or worse spatial memory impairments after severe TBI^8,36,37^ consistent with our initial hypothesis, however it should be noted that these severe open head TBI models are different pathologically from closed head mTBI and rmTBI models^38,39^.

In contrast to the spatial memory domain, anxiety-related behavior exhibited an age-dependency in this study where rmTBI increased thigmotaxis in aged animals, but not in young animals. We base this interpretation specifically on our analyses that test age-dependence rather than simply a within-age significance contrast^16^. In our Tier 2 analysis of sham-normalized rmTBI performances in EPM, percentage of time spent in the open arms (*p*=0.0140, *d*=1.12, **Fig 3D**) and closed arms (*p*=0.0208, *d*= −1.04, **Fig 3B**) differed significantly between age groups with aged animals spending less time in the open arms and more time in the closed arms compared to young. Additionally, our Tier 3 analysis showed the Age x Injury interaction effect trended in the same direction for both open arms (*p*=0.082, ω²p=0.038) and closed arms (*p*=0.078, ω²p=0.044) with no significant injury main effect but significant age main effect for both open (*p*=0.0050, ω²p=0.136) and closed arms (*p*=0.0497, ω²p=0.060), suggesting that the response is carried by a single age group rather than both. Supporting our interpretation is our Tier 1 within-age analysis showing the aged rmTBI group spent significantly less time in the open arms relative to aged sham (*p*=0.0265, *d*= −1.02, **Fig 3C**) while the young groups were unaffected.

Convergently, OFT is concordant with EPM in directionality; rmTBI increases thigmotaxis with a significant injury main effect in time spent in the outer zone (*p*=0.0207, ω²p=0.062). While a medium-large effect size was detected in both young and aged animals, only the aged groups reached within-age significance where the aged rmTBI group spent significantly more time in the outer zone (*p*=0.0468, *d*=0.89, **Fig 3E**) compared to aged shams. Since the Age x Injury interaction effect and our normalized inter-age comparisons were null, we interpret our OFT result as an rmTBI effect that is more readily resolved against an aged baseline. EPM strongly supports an age-dependent anxiety effect, with OFT corroborating the direction of the aged signal.

This age-dependent anxiety phenotype is the central novel finding of this study. Our previously published studies using this model of rmTBI in young-adult mice have shown that rmTBI reduces anxiety-like behavior and increases risk-taking behavior up to 6 mpi^17,19^. Our foundational study establishing this model of rmTBI showed mice with rmTBI spent more time in, traveled further within, and made more entries into the open arms of the elevated plus maze compared to sham and single mTBI groups suggesting a disinhibited, risk-taking profile rather than an anxious profile^17^. In a follow-up study, we reproduced this result showing rmTBI increased time spent in open arms in a dimly-lit room (5 lux white light), as well as showed rmTBI also decreased inner zone time in OFT^19^. Outside of our lab, Islam *et al.* showed similar results in a CCI model comparing young and aged mice^8^. In a well-lit Elevated Zero Maze, young CCI mice showed disinhibited, increased risk-taking behavior spending more time in the open compared to young sham and aged CCI mice while aged CCI mice were not different than aged sham mice^8^. Further, multiple studies have shown in OFT, aged CCI mice spent significantly less time in the inner zone of the OFT compared to aged sham, while young CCI mice were not statistically different than young sham^7,8^. While both EPM and OFT are both used to test anxiety-like behavior, EPM can be used to test either anxiety-related behavior or risk-taking behavior depending on room illumination^40–42^. In a well-lit or dimly-lit environment (>1 lux), open arms are aversive due to height, openness, and light, thus baseline anxiety and thigmotaxis is higher in tested animals due to multiple anxiogenic stimuli creating more dynamic range to study changes in disinhibition rather than anxiety^40^. In dark environments (<1 lux), open arms no longer have light as an anxiogenic stimuli, thus tested animals are more exploratory, creating more dynamic range to study changes in anxiety-related behavior^40,42,43^. In this study, we were primarily interested in anxiety-related behavior based on previous literature showing aged mice with CCI exhibited more anxiety-related behavior^7,8^ so we performed our EPM tests in a dark room with dim red track lighting overhead that mice are essentially blind to. In the current study, the young animals show a similar pattern to our past studies with slightly more time spent in the open arms of the EPM, but the difference between sham and rmTBI were not significant. However, our aged animals show the inverse: reduced open-arm occupancy and increased thigmotaxis, the main signature of heightened anxiety. The emergence of the large aged anxiety phenotype under these anxiety-sensitive conditions strengthens our confidence in the results. Conversely, the null result in our young animals is interpreted as an absence of an anxiety-related phenotype, supported by the OFT, rather than the absence of any affective phenotype.

Methodologically, this study illustrates that analytical strategy is inseparable from study design in factorial experiments with multiple independent variables. The direct test of whether age affects outcomes after rmTBI is the Age x Injury interaction effect. In our study, we showed significant anxiety-related deficits in the within-age Tier 1 analysis with large effect sizes. Despite this, the same anxiety-related endpoints showed small effect sizes with trending significance in the Tier 3 Age x Injury interaction. We performed a post-hoc sensitivity analysis to compute an appropriate sample size to detect a significant interaction effect in EPM, which had the largest interaction effect sizes (ω²p=0.038-0.044), with 80% power. Our post-hoc sensitivity analysis revealed that sufficient sample size to detect this interaction is n=44 animals per group (N=176 animals total). After including 10% extra animals for attrition, an Age x Injury behavioral study requires N=194 animals total, which is infeasible financially, logistically, and ethically for most single research labs. Further, this sensitivity analysis is only appropriate for testing Age x Injury interactions in one sex. To probe Age x Sex x Injury would increase the required sample size further. This is a consistent and well-recognized difficulty in powering studies to detect interaction effects relative to main effects in neuroscience^14,15^. It is tempting to treat within-age patterns as demonstrating age-dependence. If an effect is significant in one age group and null in the other, this does not demonstrate age-dependence^16^. We therefore anchor our age-dependence claims on the analyses that compare the cohorts directly, a trending interaction (Tier 3) and a significant normalized inter-age comparison (Tier 2). We acknowledge that the normalized the rmTBI groups to a fixed within-age sham mean does not propagate the variance in the sham groups, thus we view this analysis as complementary rather than a definitive substitute for the interaction term. Further, we treat the within-age (Tier 1) comparisons as concordant support. With this framing, our three tier statistical approach is not a device used to rescue null factorial results, but rather a disciplined way of separating what the data can support (convergent, direction-consistent, large-effect evidence for age-dependent anxiety) from what they cannot (a significant interaction at this sample size) and reporting that distinction transparently.

Clinically, 18-month C57BL/6 mice approximate humans of roughly 60-70 years^44,45^. Chronic anxiety and affective dysregulation are recognized, but often under-prioritzed, chronic sequelae of mTBI and rmTBI in older adults^28,46–48^. Our finding that aged animals show age-dependent anxiety-related behaviors after rmTBI in the absence of any locomotor impairment is important clinically. Post-injury monitoring of TBI often centers on physical and motor recovery. Since our aged affective phenotype arose without locomotor impairment, the practical implication is that affective sequelae may emerge in older TBI patients who otherwise appear physically recovered and could therefore be overlooked. More broadly, the domain divergence pattern shown here with shared cognitive vulnerability, but age-gated affective vulnerability, argues against our initial hypothesis that age would simply worsen rmTBI outcomes. It is more in favor of domain-specific investigations of aged brain vulnerability to injury.

## Limitations

This study has several limitations. First, only male mice were used in this study. Multiple studies have shown that there are sex-specific differences in the injury response after TBI^18,19,26,27,48–52^. Female cohorts and age x sex x injury studies are a priority for follow-up. Second, this study only includes a single chronic behavioral timepoint. A single timepoint leaves the trajectory of the observed phenotypes uncharacterized. Several studies report chronic behavioral deficits beyond 4 months, up to 24 months post-TBI^17,31,32,34^. Future studies should incorporate longitudinal behavioral testing to map trajectories of behavioral phenotypes with consideration for carryover effects as possible confounds^53^. Multiple behavioral tests for a single domain may be used to reduce carryover from animals learning the specific tasks. Third, our study includes EPM data from mice that were tested in the dark to increase sensitivity to anxiety-specific behavioral changes. However, this paradigm is relatively insensitive to disinhibition which has been reported in young adults. Therefore, the null affective phenotype in our young cohorts should be interpreted specifically as a null anxiety phenotype rather than a broad affective phenotype. Last, findings in any study utilizing a single inbred strain of laboratory rodents may not generalize across genetic backgrounds. Further, our samples sizes in this study are adequately powered for the primary within-age comparisons and main effects but underpowered for interaction detection. This limitation is precisely why we utilized a three-tiered statistical framework.

## Conclusion

In aged male C57BL/6 mice, chronic rmTBI induces a domain-divergent behavioral phenotype: shared spatial memory deficits, but age-dependent anxiety-like behavior. While this result is not supported by a factorial interaction term at typical preclinical sample sizes, it is directionally consistent and resolved by within-age and normalization-based analyses. Our findings show that age shapes the anxiety-related, but not the cognitive, consequences of chronic rmTBI, while highlighting that in studies probing age-related, or to a wider extent sex-related, effects on injury, statistical methodology is inseparable from study design. This behavioral study sets up future studies to identify cellular and molecular mechanisms that underpin this age-gated anxiety phenotype.

## Acknowledgements

We want to thank technical staff at the Sue & Bill Gross Stem Cell Center, especially Rebecca Nishi, Katelyn Anderson, and Stephanie Hernandez for their help with animal studies. We would like to thank Mathew Blurton-Jones for sharing his Barnes Maze apparatus. We would like to thank the UCI Behavioral Testing Core for assisting us in performing Open Field Test.

## Funding Declaration

This study was supported by CIRM DISC0-14447. Josh Karam was supported by the Neurobiology of Aging T32 grant. Leslie Ortiz was a CIRM intern funded through the CIRM Grant EDUC2-12638.

## Competing Interests

The authors declare that they have no competing interests.

## Ethics Declaration

All experiments, including animal housing conditions, surgical procedures, and postoperative care, were in accordance with the Institutional Animal Care and Use Committee guidelines at the University of California–Irvine (UCI).

## Notes

### Competing Interest Statement

The authors have declared no competing interest.

